# Single-cell epigenetic profiling reveals an interferon response-high program associated with *BAP1* deficiency in kidney cancer

**DOI:** 10.1101/2024.11.15.623837

**Authors:** Sabrina Y. Camp, Meng Xiao He, Michael S. Cuoco, Eddy Saad, Erica Pimenta, Kevin Meli, Ziad Bakouny, Chris Labaki, Breanna M. Titchen, Yun Jee Kang, Jack Horst, Rachel Trowbridge, Erin Shannon, Karla Helvie, Aaron R. Thorner, Sébastien Vigneau, Angie Mayorga, Jahnavi Kodali, Hannah Lachmayr, Meredith Bemus, Jihye Park, Toni Choueiri, Kevin Bi, Eliezer M. Van Allen

**Author notes:** Co-corresponding authors: Kevin Bi, Dana-Farber Cancer Institute 450 Brookline Ave, Boston MA 02215, Eliezer M. Van Allen, Dana-Farber Cancer Institute 450 Brookline Ave, Boston MA 02215.

## Abstract

Renal cell carcinoma (RCC) is characterized by recurrent somatic mutations in epigenetic regulators, which stratify patients into clinically significant subgroups with distinct prognoses and treatment responses. However, the cell type-specific epigenetic landscape of RCC—broadly and in the context of these mutations—is incompletely understood. To investigate these open questions, we integrated single nucleus ATAC sequencing data from RCC tumors across four independent cohorts. In clear cell RCC tumors, we identified four shared malignant epigenetic programs related to angiogenesis, proximal tubule-like features, interferon (IFN) signaling, and one that lacked distinct genomic regions with increased accessibility. Among the mutated epigenetic regulators, *BAP1* mutation exhibited the most significant impact on chromatin accessibility in tumor cells, and the associated epigenetic changes were linked to IFN response. We identify multiple potential sources of elevated IFN signaling in these lesions, such as increased immune infiltration and increased accessibility and expression of an IFN-associated ERV, ERV3-16A3_LTR. We find that the expression of ERV3-16A3_LTR may itself be a negative prognostic biomarker in ccRCC. Our findings highlight the convergence of malignant epigenetic programs across ccRCC tumors and suggest that *BAP1* loss, potentially through ERV3-16A3_LTR dysregulation, is associated with an IFN response-high epigenetic program.

## INTRODUCTION

Renal cell carcinoma (RCC) is a cancer partly defined by recurrent mutations in epigenetic regulators. Deficiency in these epigenetic regulators has been associated with distinct evolutionary paths, influencing core disease biology, prognosis, and therapeutic outcomes ^1–5^. For example, mutations in *BAP1* define a particularly aggressive subset of disease characterized by high tumor grade, sarcomatoid differentiation, and tumor necrosis ^6–10^. On the other hand, the near mutually exclusive *PBRM1*-deficient RCC tends to be of lower grade and may serve as a predictive biomarker of response to anti-angiogenic and immunotherapy treatments in select clinical contexts ^5,11–15^.

Although mutated epigenetic modifiers are a key feature of this cancer, prior work exploring the epigenetic landscape of RCC broadly and in the context of these mutations is limited. A recent study using bulk ATAC sequencing to profile clear cell RCC (ccRCC) identified two epigenetic programs pointing to heterogeneity in hypoxic response and immune response ^16^. However, since these findings were based on bulk accessibility profiles, the cellular origin of these programs is unknown. Another study began to dissect cell type-specific epigenetic changes associated with mutations in chromatin modifiers, *PBRM1* and *BAP1,* using single nucleus ATAC sequencing (snATAC-seq) data ^17^. However, broader conclusions from this work are limited by the sample size and clinical heterogeneity of the cohort.

To elucidate the epigenetic landscape across RCC tumors and evaluate how this landscape is impacted by the frequently mutated epigenetic regulators that characterize this disease, we integrated snATAC-seq data from RCC tumors across four independent cohorts. By deconvolving cell type from accessibility measurements, we hypothesized that we could uncover malignant epigenetic programs that converge on central aspects of RCC malignancy. Additionally, we proposed that our multicohort integrated dataset would allow us to interrogate the impact of chromatin modifier mutations on the accessibility landscape and the composition of the immune microenvironment, while accounting for clinical covariates that have confounded previous studies.

## RESULTS

### Integrating multicohort snATAC-seq data to explore the epigenetic landscape of RCC

Given that (i) across histologic subtypes, RCC has a high prevalence of mutations in epigenetic modifiers, and (ii) these mutations have been associated with differential prognoses and therapeutic responses, we first sought to characterize the cell type-specific epigenetics of RCC broadly and in the presence of certain epigenetic modifier mutations using snATAC-seq ^18^. To do so, we combined snATAC-seq data from four different cohorts, one from internally generated data and three from previously published studies ^17,19,20^. In total, our combined cohort had snATAC-seq data for 61 unique biopsies from 58 patients with RCC (Figure 1A). For our internal dataset, we generated snATAC-seq data from 16 unique tumor biopsies (n = 13 patients), all derived from frozen tissue. Four patients from this internal cohort also had additional snATAC-seq data generated using multiome single-nucleus RNA and ATAC sequencing (snRNA-ATAC-seq). The three external RCC snATAC-seq datasets were derived from a mixture of frozen and fresh samples. Most specimens across internal and external cohorts were derived from the kidney, with a minority from bone, lymph node, and visceral metastases. Two specimens had non-clear-cell subtypes, papillary and *TFE3*-translocation, while all other specimens were of clear cell histology. The combined cohort contained a mixture of disease stages and grades. Genomically, the cohort had a high frequency of somatic mutations in *VHL* and chromatin modifier genes, recapitulating known genomic characteristics of RCC (Figure 1B, Table S1)^18^. As expected, most putative tumor cells from clear cell lesions had chromosome 3p loss, along with other arm level gains and losses, compared to non-malignant cells (Figure 1C, Table S1).

**Figure 1.**
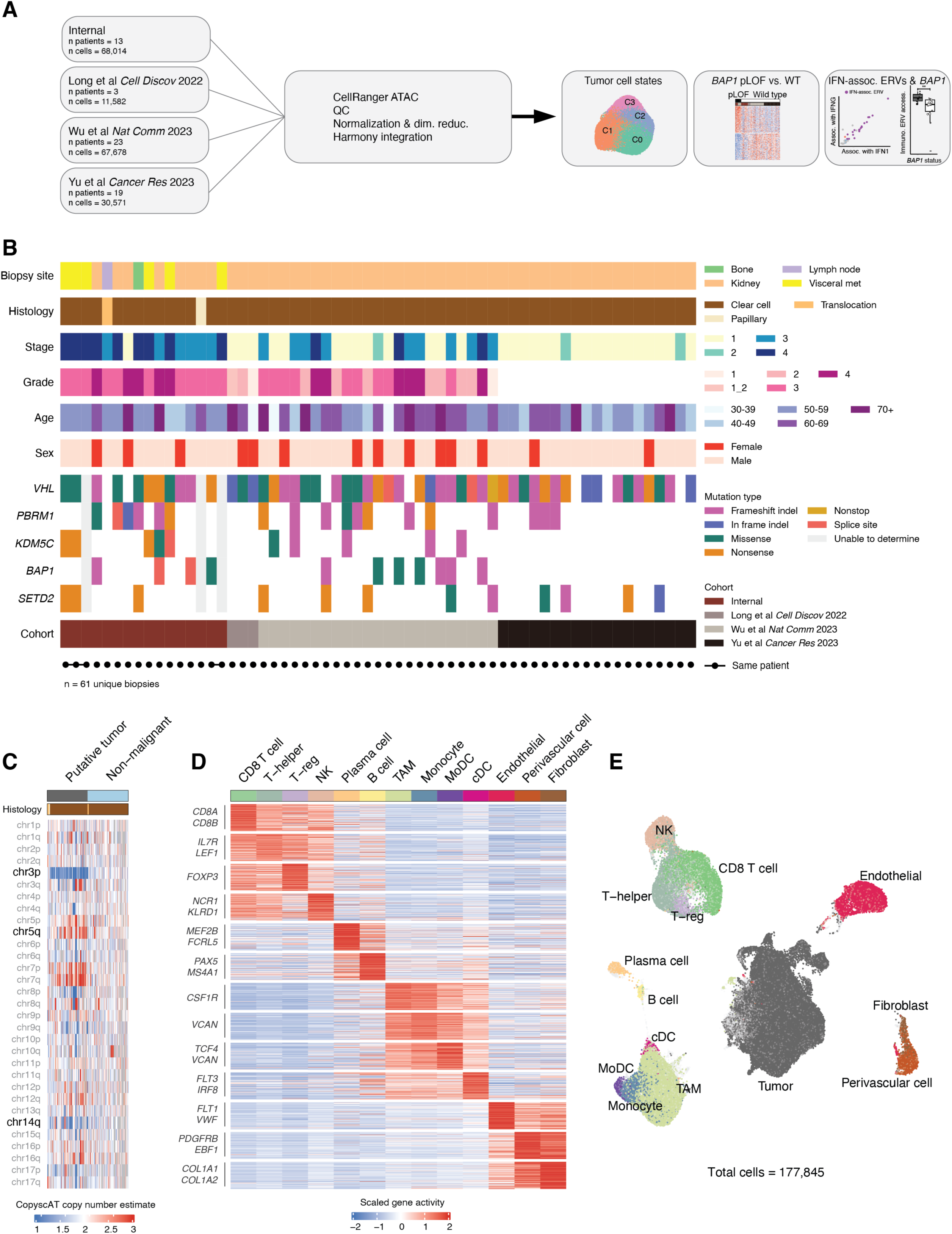
Multicohort snATAC-seq analysis of the epigenetics of renal cancer. A) Study overview B) Comutation plot of the clinical and genomic characteristics of the unique biopsies included in the study. Each row is a feature (clinical variable or gene) and each column corresponds to an individual biopsy. Translocation RCC sample identified via positive *TFE3* FISH test. C) Heatmap of a subset of chromosome arm copy number estimates derived from CopyscAT, split by putative tumor and non-malignant cells. Displaying 10,000 cells for each identity ^48^. Histology color mapping detailed in the legend for panel B. D) Heatmap of scaled average gene activity scores across non-malignant cell types, featuring top 100 differentially accessible genes per cell type based on average log2 fold-change from logistic regression adjusting for sequencing depth (nCount_ATAC, adjusted p-value < 0.05, Bonferroni correction), with logistic regression applied to downsampled identities (1000 cells per identity) and known cell type marker genes annotated within top 100 gene sets. Legend for histology annotations from panel B. E) Visualization of UMAP dimension reduction (computed on LSI components 2 through 50) for all cells included in the study. Broad cell types are annotated, and cells depicted in light grey were excluded from the analyses.

After quality control (QC), our combined dataset had a total of 177,845 cells (Methods). QC metric distributions of QC-passing cells did not vary substantially based on the originating cohort or whether the sample was fresh or frozen (Figure S1A). To detect cell types and states shared across samples and control for sample-specific batch effects, we used Harmony to adjust low-dimensional embeddings using sample ID as the batch covariate ^21^. Cell types were annotated through iterative re-clustering, evaluation of differentially accessible genes, and canonical marker gene accessibility (Figure 1D, 1E, Table S1, Methods). The cell types captured in our snATAC-seq dataset were comparable across different cohorts and sample preparation methods (fresh or frozen), allowing us to further analyze these samples in aggregate (Figure S1B).

### Characterization of chromatin accessibility in RCC

To detect distinct tumor epigenetic states shared across patient tumors, we first performed dimension reduction and unsupervised clustering within all malignant cells identified via arm-level copy number alterations. From this, we found that malignant cells predominantly clustered according to sample and/or RCC subtype. For example, we identified one cluster, annotated *RADIL*-high, that was specific to the one papillary RCC sample. This cluster had significantly increased accessibility over genes such as *RADIL*, *PGAP3*, and *HOXB7*. We also identified one other RCC subtype-specific cluster, *TRIM63*-high, which was enriched for cells from the translocation RCC sample. This cluster had increased accessibility over *TRIM63*, *TIMP2*, and *PRCD*. Clusters *PDGFRA*-high, *CADM1*-high, and *NCAM1*-high were each highly sample-specific, with all samples being of clear cell histology. However, there was a set of clusters composed of cells from several different samples (the majority of which were of clear cell histology), and we named these clusters collectively “ccRCC-balanced” (Figure 2A, 2B, S2A, Table S2).

**Figure 2.**
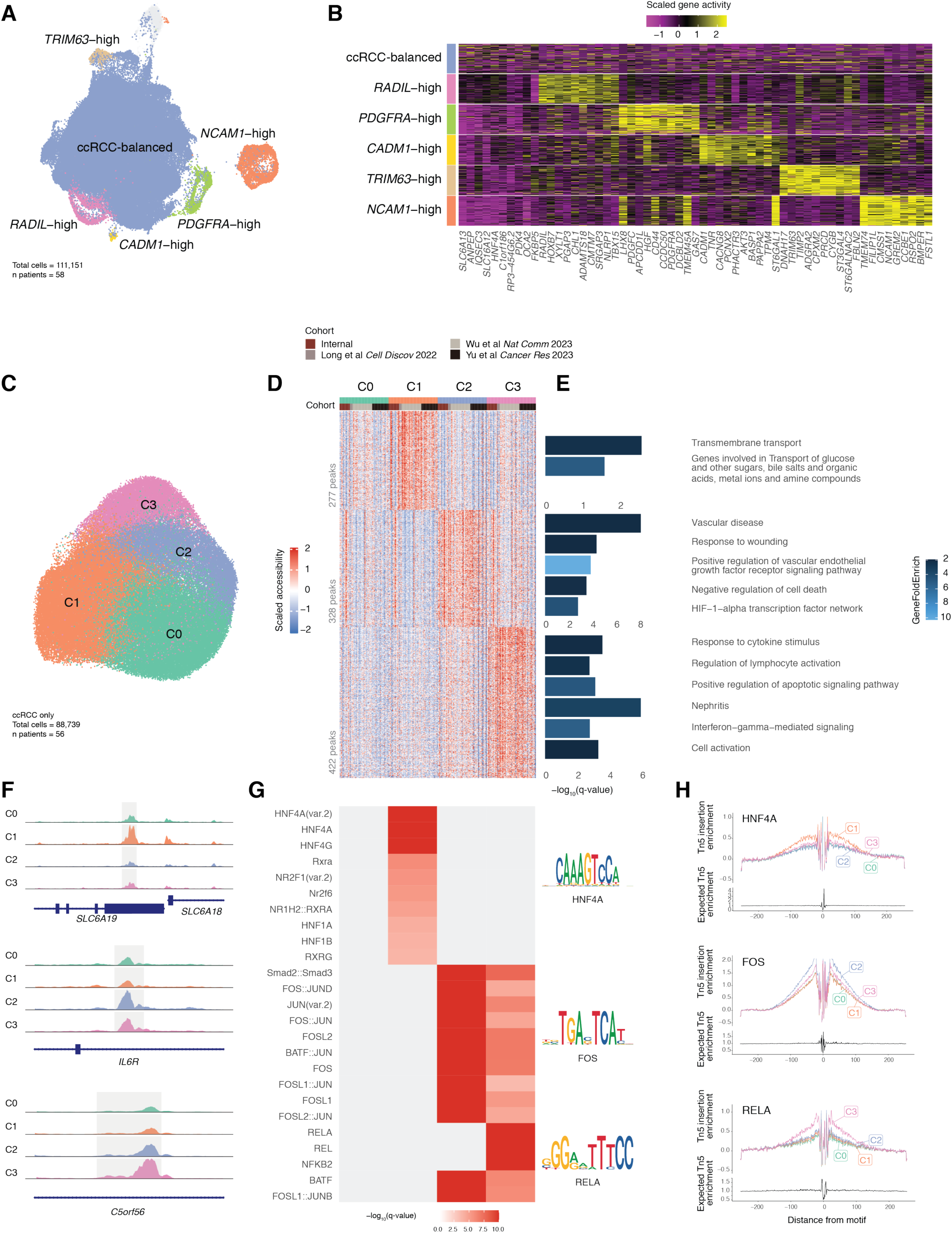
Discovery and characterization of RCC epigenetic states. A) Visualization of UMAP dimension reduction (computed on LSI components 2 through 50) of putative tumor cells from all samples, colored by malignant cell state. ccRCC-balanced denotes clusters predominantly from ccRCC lesions. Other clusters shown were enriched for specific samples/subtypes and annotated with a gene showing high accessibility in that cluster compared to all other clusters. B) Heatmap of gene activity scores across malignant cell states. Top 10 (average log2FC) differentially accessible genes for each state, identified by logistic regression (LR) adjusting for sequencing depth (nCount_ATAC, adjusted p-value < 0.05, Bonferroni correction). LR on downsampled data (1000 cells/identity), displaying 100 cells/identity. C) Visualization of UMAP dimension reduction (computed on LSI components 2 through 20) of the four largest shared clusters from ccRCC-only cells, sub-clustered from ccRCC-balanced state. Excludes highly sample-specific clusters. D) Heatmap of peak accessibility across ccRCC clusters. Columns represent scaled average peak accessibility per cells from a given sample in a particular cluster (e.g., one column is cells from sample 1 in cluster 1). Includes all significant peaks from LR (Bonferroni correction, adjusted p-value < 0.05), with columns ordered by cluster, then cohort. E) Manually selected pathways enriched in significant peaks per cluster, as determined by GREAT (Binomial test, FDR q-value < 0.05). GeneFoldEnrich indicates fold enrichment of the number of genes in the test set with the set of genes in the given pathway. Pathways selected from GO biological process, disease ontology, MSigDB pathway ontologies. F) Coverage plots for selected peak-gene associations identified by GREAT that were included in significant pathways shown in Figure 2E. Coverage plots split and colored by ccRCC cluster. G) Heatmap of -log10 q-values (hypergeometric test, FDR q-value < 0.05) for TF motif enrichment in significant peak sets. Cutoff of -log10 q-value at 10. Top 10 motifs per cluster shown; position weight matrices for HNF4A, FOS, RELA to the right. H) Transcription factor footprinting plots shown for HNF4A, FOS, and RELA. Footprinting tracks split and colored by ccRCC cluster

### Dissecting tumor cell states in ccRCC

To more deeply discern epigenetic variation within this shared tumor state, we sub- clustered ccRCC-balanced with only cells from ccRCC biopsies (n = 56 tumors; 88,739 tumor cells). We identified four tumor states shared across patients, cohorts, and disease stages, and annotated them as C0, C1, C2, and C3 (Figure 2C, S2B). To nominate genomic regions whose accessibility defined each state, we performed differential peak accessibility analysis (Methods). We found 1,027 regions to have significantly different accessibility across these states. There were no peaks enriched for accessibility in the C0 state, suggesting that this state may be more epigenetically quiescent compared to the other states (Figure 2D, Table S2).

To nominate biological properties reflected by each state, we performed pathway enrichment analysis on state-specific genomic regions using GREAT ^22^. We found that peaks uniquely accessible in tumor state C1 were involved in pathways related to kidney function, such as transmembrane transport and transport of glucose, and bile salts, among others. Peaks more accessible in tumor state C2 were attributed to pathways related to angiogenesis, such as vascular disease, HIF-1-alpha transcription factor network, and regulation of VEGF signaling. Lastly, tumor state C3 had increased accessibility over peaks involved in immune response and inflammation, such as interferon (IFN) -gamma-mediated signaling, nephritis, and response to cytokine stimulus (Figure 2E, Table S2). We also evaluated individual peaks that contributed to the observed pathway enrichments. The promoter region of *SLC6A18* had over 3-fold greater accessibility in cluster C1 compared to all other clusters and contributed to the signal in the transmembrane transport pathway. An intronic peak in *IL6R*, attributed to the vascular disease pathway, had 2-fold greater accessibility in C2, and may be generally involved in the increased angiogenesis signal in these cells. Lastly, a region downstream from *IRF1* was attributed to several immune activation and IFN response-related pathways and was enriched for accessibility in C3 (Figure 2F).

To determine which transcription factors (TFs) may be acting at the accessible chromatin regions that define these tumor states, we then tested if these peak sets were enriched for particular TF motif binding sites relative to a broadly accessible reference peak set matched for relevant genomic characteristics (Methods). Indeed, these epigenetically defined ccRCC tumor states had uniquely active TFs. C1 peaks were enriched for binding site motifs of TFs related to kidney function, such as HNF4A, a key regulator of proximal tubule development ^23^. C2 peaks were enriched for FOS and JUN TF motifs, among others. Lastly, the regions with increased accessibility in the C3 program were enriched for motifs of the NF-kB family of transcription factors (e.g., REL, RELA, NFKB2), which are closely linked to the innate immune system and type I IFN signaling (Figure 2G, Table S2). TF footprinting analysis recapitulated these results, where accessible regions across the genome that contained the selected enriched motifs had greater accessibility in tumor cells of the respective epigenetic state (Figure 2H).

### Influence of BAP1 mutation on the epigenetic landscape of ccRCC

Given that epigenetic modifier mutations have been associated with differential biological trajectories and patient outcomes, we investigated whether the accessibility profile of tumors with a particular epigenetic modifier mutation, determined at the bulk-level, was similar to one of the shared ccRCC tumor epigenetic states identified.

Biopsies with a somatic putative loss-of-function (pLOF) *BAP1* mutation had significantly greater accessibility over the immune response and inflammation-related C3 peak set relative to a reference peak set matched for relevant genomic characteristics, with the trend holding when limiting to biopsies from patients with advanced disease (Figure 3A, S3A, S3B, Methods). Although sarcomatoid differentiation has been previously associated with an inflamed tumor microenvironment (TME) and *BAP1* mutation, this feature did not appear to be the primary factor driving the observed association herein (Figure S3C) ^8,10,24^.The finding of enhanced accessibility around immune response and inflammation-related genomic regions in *BAP1*-mutant tumor cells is concordant with previous bulk transcriptomics and proteomics studies in RCC, which reported a relationship between *BAP1* mutation and IFN signaling ^24–26^. In contrast, biopsies with mutations in *PBRM1*, *SETD2*, or *KDM5C* were not enriched for accessibility over any of the shared tumor state peak sets (Figure S3A, S3B).

**Figure 3.**
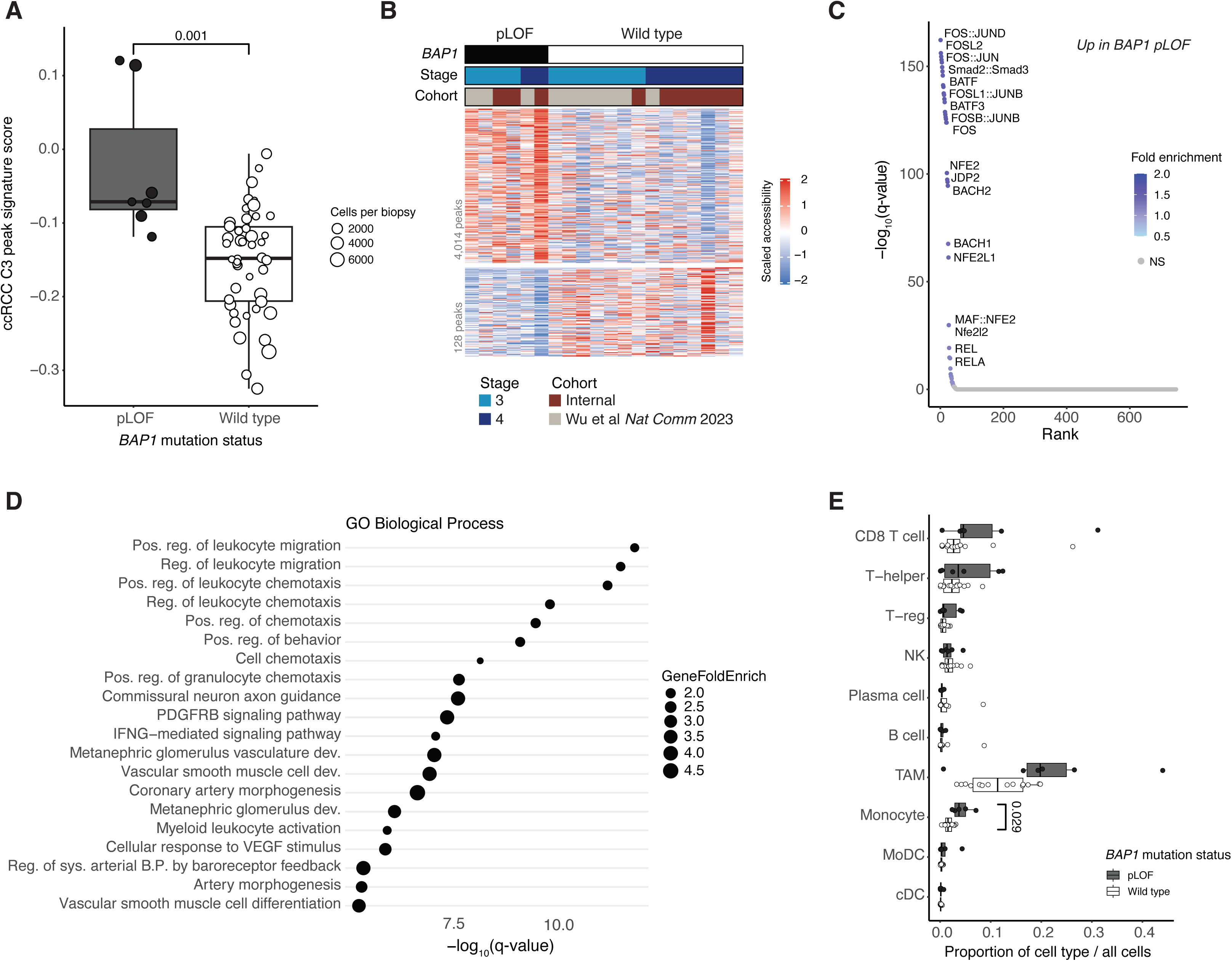
*BAP1* pLOF mutation associated epigenetic changes in advanced ccRCC. A) Boxplot of median ccRCC C3 peak signature score per unique biopsy, split by *BAP1* mutation status. *BAP1* wild type excludes pLOF and other non-silent mutations. Analysis limited to cells from ccRCC tumor states (C0,C1,C2,C3). Q-value displayed (two-sided Wilcoxen, FDR correction). B) Heatmap of scaled peak accessibility scores across *BAP1* mutation status. Analysis limited to cells from ccRCC tumor states (C0,C1,C2,C3) and from patients with advanced disease. Columns represent scaled average peak accessibility per biopsy, ordered by *BAP1* mutation status, disease stage, then cohort. Displaying all significant peaks from LR (Bonferroni correction, adjusted p-value < 0.05, log2 fold-change > abs(1)). Number of peaks relative to size of heatmap not to scale. C) Transcription factor motif enrichment in *BAP1* pLOF mutation-associated peaks identified by hypergeometric test. Annotating all significant motifs (FDR correction, q-value < 0.05), with some excluded due to label overlap. Point color indicates fold enrichment of motif presence in input peak set versus reference set. Grey points indicate q-value > 0.05. All points ordered from smallest to largest q-value left to right. D) Top 20 GO biological process terms enriched in *BAP1* pLOF significant peak set (Binomial test, FDR correction, q-value < 0.05). Dot size corresponds to GeneFoldEnrich, indicating fold enrichment of genes in test set/pathway as determined by GREAT. E) Boxplots comparing broad immune cell type proportions between biopsies with pLOF *BAP1* mutation and *BAP1* wild type, restricted to advanced stage disease patients. Points overlaying boxplots show per biopsy cell type proportions. Q-value for monocyte proportion comparison shown (two-sided Wilcoxen, FDR correction); other cell type comparisons not significant.

Consistent with prior studies, patients with *BAP1*-mutated tumors were enriched for advanced stage disease (Figure S3D) ^7,27^. Thus, to investigate *BAP1*-related epigenetic changes that were not potentially confounded by clinical disease stage, we limited all subsequent analyses to samples from patients with advanced disease. This restriction also confined our analysis to samples from two cohorts: our internal dataset and the external Wu et al. dataset. To further define the epigenetic differences associated with *BAP1* mutation, we performed a supervised differential peak accessibility analysis (Methods). We identified 4,014 regions with increased accessibility and 128 regions with depleted accessibility in tumor cells from *BAP1*-mutated lesions relative to tumor cells from *BAP1* wild type lesions (Figure 3B, Table S3). We did not find globally decreased chromatin accessibility in tumor cells from *BAP1*-mutated lesions relative to other ccRCC molecular subtypes in this multicohort analysis, in contrast to a prior study.

However, our analysis indicates that the prior finding may have been a cohort-specific observation ^17^ (Figure S3E). Consistent with the enrichment for the C3 peak set related to IFN signaling, regions with increased accessibility in tumor cells from *BAP1*-mutated lesions were enriched for binding site motifs of the NF-kB family of transcription factors, along with FOS and JUN transcription factor motifs (Figure 3C, Table S3). Peaks with depleted accessibility associated with *BAP1* mutation were not significantly enriched for any particular TF binding sites. Peaks with increased accessibility in tumor cells from *BAP1*-mutated lesions were enriched for a number of immunological processes such as regulation of leukocyte chemotaxis and IFN-gamma-mediated signaling, among others (Figure 3D, Table S3). However, peaks with decreased accessibility were not significantly enriched for any annotated pathways.

Thus, we found that tumor cells from *BAP1*-mutated ccRCC tumors exhibited a distinct epigenetic profile, characterized by increased accessibility around genomic regions corresponding to IFN response-related pathways and TF binding site motifs, epigenetically distinguishing *BAP1*-mutated ccRCC from other molecular subtypes.

### Immune infiltration in BAP1-mutated ccRCC

Next, we evaluated the potential sources of IFN in *BAP1*-mutated lesions that may contribute to the observed epigenetic profile. Prior studies have found that *BAP1* mutation is linked to increased immune infiltration, including CD8+ and CD4+ T cells, macrophages, NK cells, and others, in genetically engineered mice and patient-derived xenografts of ccRCC ^8,24^. Leveraging the single cell resolution of our ccRCC dataset, we investigated the relationship between *BAP1* mutation status and composition of the immune microenvironment. We found a significantly higher proportion of monocytes in *BAP1*-mutated tumors. Other immune cell types, specifically CD8 T cells and tumor-associated macrophages, were more prevalent in *BAP1*-mutated tumors, but these differences did not reach statistical significance (Figure 3E). Given that RCC with sarcomatoid differentiation is enriched for *BAP1* mutation and this histopathological feature has itself been associated with increased CD8+ T cell infiltration in prior studies, we considered whether these immune infiltrate associations might be influenced by sarcomatoid differentiation ^10,28^. However, our analysis did not suggest that sarcomatoid differentiation was the primary factor driving these associations in this cohort (Figure S3F).

### Identifying IFN-associated ERVs in ccRCC

While increased immune infiltration in *BAP1*-mutated lesions may serve as one, extrinsic source of IFN signaling, tumor-intrinsic mechanisms may also contribute. A previous study found *BAP1*-mutated ccRCC tumors were associated with high potentially immunogenic endogenous retrovirus (ERV) transcriptional activity ^29^. ERVs can activate a cell’s viral response machinery and lead to a viral mimicry-induced IFN response through the activation of NF-kB and IRF TFs ^30–33^. We thus hypothesized that epigenetic dysregulation of ERVs could be a mechanistic link contributing to the association between *BAP1* mutation and the IFN response-high tumor epigenetic profile observed herein.

Given the apparent cell type-specific regulation of ERVs ^34,35^ and the likely varying immunogenicity among different ERVs, we aimed to identify ERVs in ccRCC tumor cells whose accessibility is significantly associated with IFN signaling as one component of investigating the potential role of ERVs in the *BAP1* mutation-associated epigenetic profile.

To epigenetically measure IFN signaling in ccRCC tumor cells, we devised an approach that leveraged paired transcriptomic and chromatin accessibility data from our multiome snRNA-ATAC-seq samples. Specifically, we identified a set of genomic regions whose accessibility was significantly associated with transcriptomic measures of IFN1 and IFNG signaling, and used accessibility across these regions as a proxy for IFN signaling (Methods, Table S4). This approach allowed us to measure IFN signaling in our larger snATAC-seq-only dataset. In tumor cells with paired snRNA-ATAC-seq data, we observed a strong positive correlation between the normalized accessibility across these IFN-associated peak sets and the transcriptomic-based measures of IFN1 and IFNG signaling. (Figure 4A, Methods).

**Figure 4.**
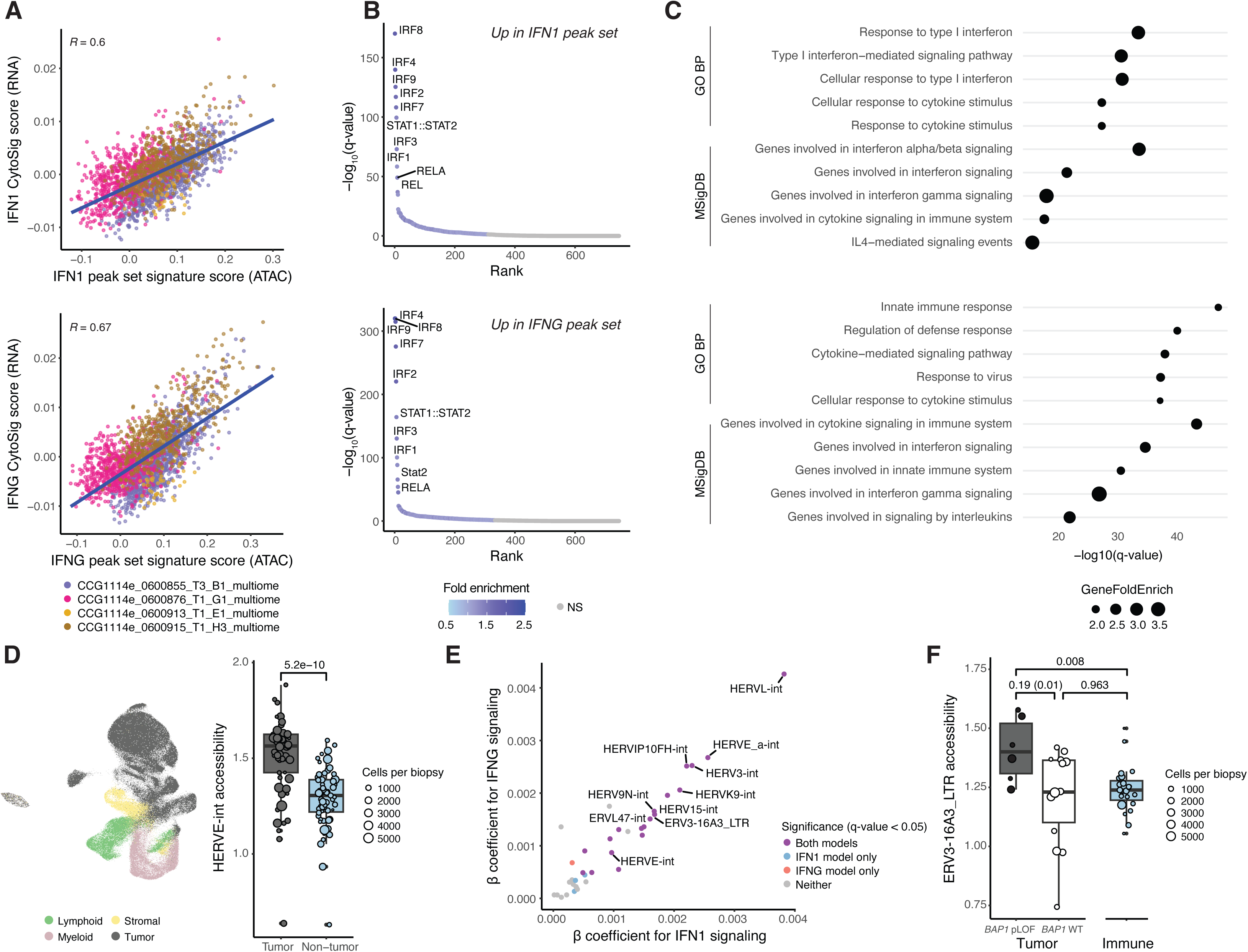
Identifying IFN-associated ERVs and dissecting their relationship to *BAP1* mutation status in ccRCC tumor cells. A) Scatter plots of RNA-derived IFN1 signaling (CytoSig) versus ATAC-derived IFN1 signaling (signature scoring for peaks associated with CytoSig IFN1 score, Methods) for all malignant cells from multiome ATAC-RNA samples (top), and IFNG comparison (bottom). Pearson correlation coefficient and regression line (blue) included; point color indicates sample ID. B) Transcription factor motifs enriched in IFN1-associated (top) and IFNG-associated (bottom) peak sets from hypergeometric test. Top 10 motifs by q-value (FDR correction) annotated; point color shows fold enrichment; grey points indicate q-value > 0.05, ordered by q-value. C) Top five GO biological process (GO BP) and MSigDB pathway annotations enriched in IFN1-associated (top) and IFNG-associated (bottom) peak sets by q-value (Binomial test, FDR correction, q-value < 0.05). Dot size corresponds to GeneFoldEnrich from GREAT. D) Left, visualization of UMAP dimension reduction (computed on PC components 1 through 15) derived from transposable element (TE) accessibility counts of cells from ccRCC tumor biopsies that had TE quantification and passed peak-based QC. Cells colored by broad lineage designation determined from peak-based analysis. Right, boxplot comparing median HERVE.int accessibility counts in a given biopsy’s putative tumor cells to non-tumor cells. Size of dot corresponds to the number of tumor cells or non-tumor cells captured in each biopsy. Nominal p-value determined from one-sided Wilcoxen test. E) Scatter plot of mixed effects model coefficients for individual ERVs from the epigenetic IFN1 signaling outcome model (x-axis) and epigenetic IFNG signaling outcome model (y-axis), showing ERVs with estimates > 0. Point color denotes significance and comparison (FDR correction); annotated ERVs significantly associated with both IFN1 and IFNG peak set signature scores (q-value < 0.05, coefficient > 0). F) Boxplot comparing the median accessibility of ERV3-16A3_LTR in ccRCC tumor cells with pLOF *BAP1* mutations, *BAP1* wild-type cells, and immune cells from biopsies with known *BAP1* mutation status. Immune cells are defined as cells from the myeloid or lymphoid lineage. Only stage 3 and 4 biopsies are included. Dot size represents the number of cells per biopsy. Q-value shown for *BAP1* mt vs. *BAP1* wt comparison (one-sided Wilcoxon test, FDR correction) and nominal p-value (one-sided Wilcoxen test) shown in parentheses; P-values (two-sided Wilcoxon test) shown for *BAP1* mt vs immune and *BAP1* wt vs immune comparison

To further confirm that the IFN1 and IFNG peak sets were an accurate reflection of IFN signaling, we evaluated which TF binding site motifs were enriched in each peak set relative to a reference set of genomic regions matched for relevant genomic characteristics. We found that both IFN1 and IFNG peak sets were significantly enriched for binding site motifs of IFN-related TFs such as IRFs, STATs, REL, and RELA (Figure 4B, Table S4). For each peak set, we also performed a pathway enrichment analysis. The set of regions whose accessibility was associated with IFN1 signaling were attributed to pathways related to type I IFN signaling and response to cytokine stimulus. The set of regions whose accessibility was associated with IFNG signaling were enriched for similar annotations, such as pathways related to immune response, IFN gamma signaling, and response to cytokine stimulus (Figure 4C, Table S4).

To measure the accessibility of ERVs at a single cell level, we used scTE, a single-cell transposable element (TE) quantification tool ^34^. First, we evaluated scTE’s performance in our dataset by testing whether TE accessibility measurements alone could distinguish broad cell types, a key finding from their study. In our ccRCC snATAC-seq dataset, these measurements were indeed able to differentiate broad cell lineages. Furthermore, we investigated the tumor-specific activity of HERVE.int in ccRCC and found supporting evidence for its activity using chromatin accessibility data, complementing previous findings based on RNA expression ^36^(Figure 4D).

With the per cell measurements of IFN signaling and ERV accessibility, we identified IFN-associated ERVs in tumor cells by testing whether the accessibility of each ERV was significantly associated with our epigenetic measure of IFN signaling, controlling for relevant covariates in a mixed effects model (Methods). We identified 19 ERVs that were significantly associated with both higher IFN1 and IFNG epigenetic scores (Figure 4E, Table S4). This set of IFN-associated ERVs included HERVE.int, an ERV that has been shown to produce a highly immunogenic tumor antigen in ccRCC ^36,37^.

### BAP1 mutation status and IFN-associated ERVs

Evaluating how these IFN-associated ERVs were related to *BAP1* mutation status, we found that the accessibility of ERV3-16A3_LTR was nominally significantly greater in tumor cells from *BAP1*-mutant lesions compared to tumor cells from *BAP1*-wildtype (Figure 4F, S4A). This ERV dysregulation appeared to be specifically *BAP1* mutation related rather than broadly associated with tumor cells; for example, there was no difference in accessibility between immune cells and *BAP1*-wildtype tumor cells (Figure 4F). Prior literature surrounding this ERV focused on the RNA expression of the internal region, ERV3-16A3_I. Such research has demonstrated that dsRNA derived from ERV3-16A3_I robustly activates a cell’s viral response machinery via the RIG-I pathway. This activation is indicated by the increased mRNA and protein expression of RIG-I-like receptors and type I IFNs in CD4+ T cells from lupus patients ^38^. Additionally, ERV3-16A3_I is a component of the MHC class I region lncRNA *HCP5*, which has been implicated in both adaptive and innate immune responses ^39^.

While increased accessibility at this ERV in malignant cells from *BAP1*-mutant tumors may suggest a corresponding increase in the ERV’s transcription, accessibility does not always closely track with expression ^40^. To investigate the relationship between ERV3-16A3_LTR expression and *BAP1* mutation in ccRCC, we quantified ERV expression in bulk RNA sequencing (RNA-seq) data from the TCGA clear cell renal cell carcinoma (KIRC) cohort using scTE (Methods). We found that expression of ERV3-16A3_LTR was significantly higher in *BAP1*-mutated tumors compared to *BAP1*-wildtype, as determined by a linear model controlling for tumor purity and clinical disease stage among other relevant covariates (Methods, Figure 5A). When stratifying the analysis by disease stage, we observed that this association was stronger in patients with early-stage disease (Figure S5A, S5B).

**Figure 5.**
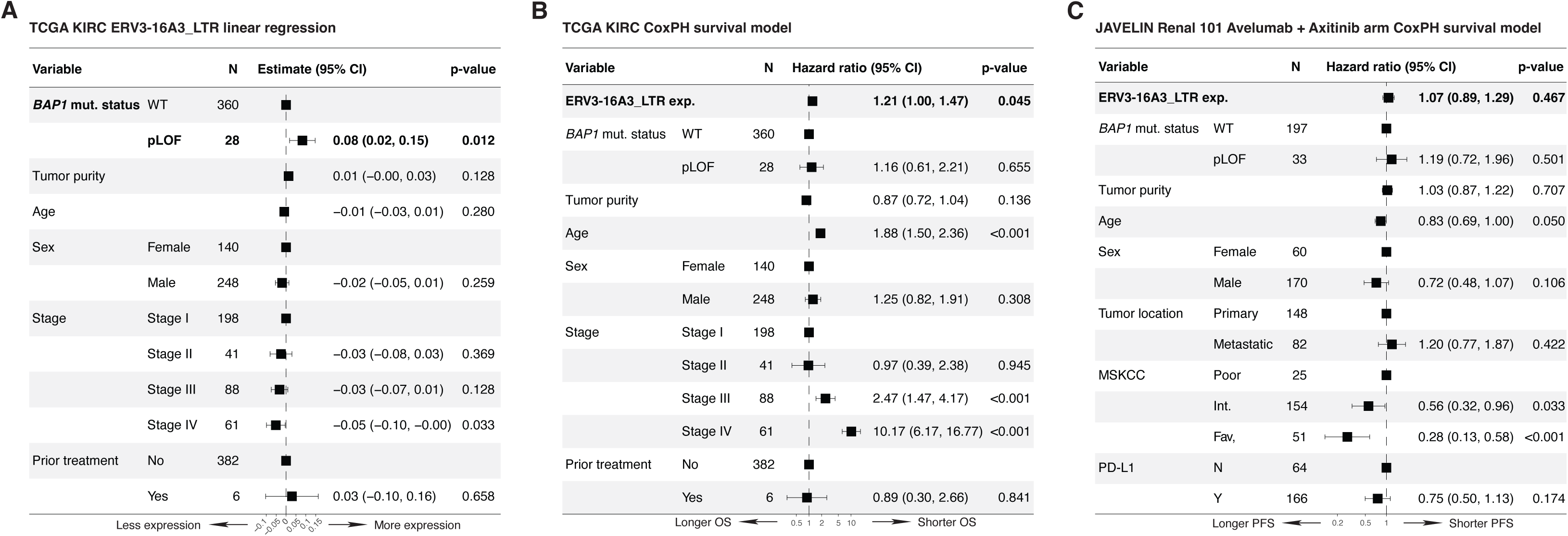
Evaluating relationship between ERV3-16A3_LTR expression, *BAP1* mutation status, and clinical outcomes in bulk ccRCC RNA-seq cohorts. A) Forest plot showing beta coefficients from a linear regression model for expression of ERV3-16A3_LTR in TCGA KIRC cohort B) Forest plot showing hazard ratios from a Cox proportional hazards model for overall survival in TCGA KIRC cohort C) Forest plot showing hazard ratios from a Cox proportional hazards model for progression-free survival in ICB-containing treatment arm of JAVELIN Renal 101 cohort

### ERV3-16A3_LTR expression and clinical outcomes in RCC

Given the evidence herein that ERV3-16A3_LTR activity is associated with IFN signaling in tumor cells, which has been associated with tumor-mediated immune escape in melanoma ^41^, we hypothesized that ERV3-16A3_LTR expression could have prognostic relevance beyond its relationship to *BAP1* mutation status in ccRCC. We found that higher expression of ERV3-16A3_LTR was significantly associated with worse overall survival in the TCGA KIRC cohort, even after controlling for *BAP1* mutation status and other covariates such as clinical disease stage, age, and tumor purity (Methods, Figure 5B). When stratifying the analysis by disease stage, we observed that this association was stronger in patients with early-stage disease (Figure S5C, S5D).

To ask whether ERV3-16A3_LTR expression may predict clinical outcomes under immunotherapy, perhaps as an indicator of tumor-mediated immune escape mechanisms, we utilized pre-treatment bulk RNA-seq data from JAVELIN Renal 101, a large cohort of RCC patients with advanced stage disease treated with first-line immune checkpoint blockade (ICB) plus anti-VEGF combination or single-agent sunitinib ^42,43^.

However, we did not find that ERV3-16A3_LTR expression was significantly predictive of response or resistance to ICB or non-ICB containing regimens (Methods, Figure 5C, S5E). Additional factors relevant to ICB outcomes in RCC, such as TME-related variables, may need to be incorporated into the model to better understand if there is a relationship between ERV3-16A3_LTR expression and current standard-of-care ICB/anti-VEGF outcomes.

Although the relationship between *BAP1* mutation and ERV3-16A3_LTR expression in TCGA KIRC appeared to be strongest in tumors from patients with early stage disease, we lastly evaluated this relationship in JAVELIN Renal 101, which consists of tumors from patients with advanced disease only. We did not find supporting evidence of greater ERV3-16A3_LTR expression with *BAP1* mutation in JAVELIN Renal 101, suggesting stage-specific biology may play a role in this association (Figure S5F).

Thus, we found that in ccRCC tumor cells, there are several ERVs whose chromatin accessibility is associated with IFN signaling. One of these IFN-associated ERVs, ERV3-16A3_LTR, is more accessible and expressed with *BAP1* deficiency, with stage-specific biology potentially playing a role in these associations. The dysregulation of this ERV may contribute to the observed phenotype herein of a *BAP1* mutation-associated IFN response-high epigenetic profile, by promoting a viral mimicry response in tumor cells. Expression of ERV3-16A3_LTR may be an independent negative prognostic biomarker in ccRCC, although the mechanism underlying the ERV’s association to distinct clinical outcomes requires further research.

## DISCUSSION

To explore the epigenetic landscape of RCC—a cancer characterized by frequent mutations in chromatin-modifying genes—we assembled a multicohort snATAC-seq dataset. Broadly, chromatin accessibility profiles of malignant cells varied significantly between RCC subtypes. Within the ccRCC subtype, we identified four major epigenetic programs that are detected in nearly all patients’ tumors. These programs describe known axes of variation in ccRCC such as angiogenesis, proximal tubule-like features, and IFN stimulated programs.

Mutations in the most frequently altered epigenetic regulators in RCC, such as *PBRM1*, *SETD2*, and *KDM5C*, did not associate significantly with any of the identified ccRCC epigenetic programs. More samples may be needed to detect more subtle chromatin accessibility changes induced by these mutated modifiers. By contrast, the accessibility profiles of *BAP1*-mutated tumor cells were significantly enriched for an IFN response-high epigenetic program, a relationship supported by prior work in other data modalities ^24–26^. Consistent with this program enrichment, chromatin with increased accessibility in *BAP1*-mutant tumor cells were involved in immunological processes and enriched for IFN-related TF binding site motifs.

We find evidence for both tumor extrinsic and intrinsic mechanisms of IFN signaling present in *BAP1* mutated tumors that may contribute to the associated epigenetic profile. In our dataset and in line with prior in vivo studies of RCC ^8,24^, *BAP1* mutated tumors showed a higher frequency of infiltrating CD8+ T cells, which are known to be a major source of IFNG in the tumor microenvironment ^44^, although a larger dataset is needed to confirm this association. Panda et al’s work suggested a tumor intrinsic source of IFN signaling in *BAP1*-mutated ccRCC, finding that these lesions have increased immunogenic ERV expression ^29^, which can activate cytosolic sensing cascades and initiate a type I IFN response ^30–33,45^. In our dataset, we identified several ERVs whose accessibility was associated with increased IFN signaling in ccRCC tumor cells. This included HERVE.int, which has been previously reported to produce immunogenic antigens in ccRCC ^36^.

The IFN-associated ERV ERV3-16A3_LTR was significantly more accessible and expressed with *BAP1* mutation, with stage-specific biology potentially playing a role in the association. Prior work supports ERV3-16A3_LTR’s relationship with IFN response, where it has been shown that dsRNA derived from the internal region of this ERV element can strongly trigger a cell’s innate antiviral immune response ^38^. Future research should aim to identify if specific ERV3-16A3_LTR loci are driving this signal to narrow down the experimental search space. If those are identified, it will be critical to determine the dependency of the *BAP1* associated phenotype on this ERV element by perturbing ERV expression in a *BAP1*-deficient ccRCC cell line and evaluating the effects on IFN signaling.

Prior work has connected IFN signaling in tumor cells to tumor-mediated immune escape. For example, in melanoma mouse models, one cancer cell IFN-stimulated gene, *RIPK1*, promotes tumor cell resistance to the indirect TNFRSF-mediated killing mechanism ^41^. In bulk ccRCC RNA-seq datasets, we evaluated the prognostic value of ERV3-16A3_LTR expression for overall survival and its ability to predict outcomes in ICB-treated patients. In the TCGA KIRC cohort, higher ERV3-16A3_LTR expression was associated with shorter overall survival, independent of *BAP1* mutation status, particularly in early-stage patients. In the JAVELIN Renal 101 clinical trial cohort, expression of ERV3-16A3_LTR was not significantly associated with treatment response or resistance to ICB. However, further studies incorporating TME-related factors may be necessary to fully capture this relationship, if it exists.

The observed association between *BAP1* mutation and increased immune cell infiltration, along with an IFN response-high state, presents a paradox given the aggressive nature of *BAP1*-mutant tumors. This discrepancy raises questions about the functional state of the infiltrating immune cells, particularly CD8+ T cells, and whether they are actively engaging the tumor. It also suggests the need to investigate potential mechanisms by which CD8+ T cell function is suppressed in *BAP1*-mutated lesions.

Additionally, tumor-intrinsic factors, such as increased ERV3-16A3_LTR activity as described here, might contribute to immune escape through increased IFN signaling in *BAP1*-mutated tumor cells. Elucidating these mechanisms may reveal new therapeutic targets to counteract the potential immune evasion strategies of this aggressive ccRCC molecular subtype.

## ACKNOWLEDGEMENTS

We thank the patients who participated in this study. We would like to thank Aviv Regev for her contributions and oversight in data generation for this study. This project was supported by NIH R01CA278980, R37CA222574, R50CA265182, T32GM008313, the Parker Institute for Cancer Immunotherapy, The Ambrose Monell Foundation, and the National Science Foundation GRFP DGE1144152.

## AUTHOR CONTRIBUTIONS

Conceptualization: SYC, MXH, EMVA; Data curation: ES, ZB, CL, JH, RT; Formal analysis: SYC; Funding acquisition: MXH, JP, EMVA; Investigation: MSC, KH, ART, SV, JK, HL, MB; Project administration: SYC, MXH, JH, RT, ES, AM, JP; Resources: KH, ART, SV, TC, EMVA; Supervision: KH, ART, SV, TC, KB, EMVA; Validation: SYC; Visualization: SYC; Writing -- original draft: SYC, MSC, ART, SV, KB, EMVA; Writing -- review & editing: MXH, ES, EP, KM, ZB, CL, BMT, YJK, JH, RT, ES, KH, AM, JK, HL, MB, JP, TC

## METHODS

### Internal RCC patient cohort sample collection and dissociation for snATAC-seq and multiome snRNA-ATAC-seq

Nuclei isolation was performed as previously described ^46^, using low-retention microcentrifuge tubes (Fisher Scientific, Hampton, NH, USA) throughout to minimize nuclei loss. The cohort included both OCT-embedded and snap-frozen tumor tissues. For OCT-embedded samples, an additional step involved separating the tissue from OCT by carefully excising it with tweezers and scalpels. Tissues were manually dissociated by finely chopping with spring scissors for 10 minutes, homogenized in TST solution, filtered through either a 35 µm FACS tube filter for snATAC-seq or a 30 µm MACS SmartStrainer for multiome snRNA-ATAC-seq (Miltenyi Biotec, Germany), and pelleted by centrifugation at 500g for 10 minutes at 4°C. The nuclei pellet was resuspended in a lysis buffer to permeabilize the nuclei before pelleting by centrifugation for 10 minutes at 500g at 4C. The final nuclei pellet was resuspended in 100 ul of 10x Genomics Diluted Nuclei Buffer, and trypan blue-stained nuclei were counted by eye using INCYTO C-Chip Neubauer Improved Disposable Hemacytometers (VWR International Ltd., Radnor, PA, USA).

Approximately 16,000–25,000 nuclei per sample were loaded per channel of the Chromium Next GEM Chip for processing on the 10x Chromium Controller (10x Genomics, Pleasanton, CA, USA). For snATAC-seq, transposition and library construction were carried out as per the manufacturer’s instructions (Chromium Next GEM Single Cell ATAC User Guide). For multiome snRNA-ATAC-seq, both transposition and cDNA generation, followed by library construction, were performed according to the Chromium Next GEM Single Cell Multiome ATAC + Gene Expression User Guide (Rev F). Libraries from both methods were normalized and pooled for sequencing on two NovaSeq SP-100 flow cells (Illumina, Inc., San Diego, CA, USA).

### snATAC-seq data availability

FASTQs for the internal cohort were deposited in phs002065.v2.p1. For the Wu et al cohort, FASTQs were obtained through the Cancer Data Service after dbGaP approval phs001287.v16.p6. For the Long et al and Yu et al cohort, FASTQs were obtained from the Sequence Read Archive. Cohort characteristics are summarized in Supplementary Table 1.

### snATAC-seq FASTQ processing and fragment file generation

For snATAC-seq samples and multiome snRNA-ATAC-seq samples, we used CellRanger ATAC v2.0.0 and CellRanger ARC v2.0.0, respectively, for read filtering, alignment, and cell calling. Reads were aligned to the GRCh38 human genome reference (CellRanger reference: refdata-cellranger-arc-GRCh38-2020-A-2.0.0). The corresponding fragment file and fragment file index output were used for downstream analysis.

### Quality control of snATAC-seq data

To ensure the reliability of snATAC-seq data, we implemented an extensive QC process at the individual sample level. First, we generated a filtered fragment file to only include fragments from cell-associated barcodes as determined by CellRanger’s cell calling algorithm. Using this fragment file, we identified accessible peaks using MACS2 v2.2.9.1 within Signac v1.10.0. Count matrices for each sample were constructed by quantifying insertion events within sample-specific peak sets for each cell using Signac’s FeatureMatrix function.

We identified doublets with scDblFinder^47^ and removed these cells. Subsequently, we evaluated each cell for several quality metrics: the fraction of reads in peaks (FRiP), the proportion of reads aligning to regions associated with technical/artifactual signal from ENCODE (blacklist fraction), the nucleosome signal (mononucleosome to nucleosome-free fragment ratio), transcription start site (TSS) enrichment score (TSS-centered to TSS-flanking fragment ratio), and the count of fragments overlapping peaks.

To set objective QC thresholds, we used a statistical approach based on the median and median absolute deviation (MAD) calculated on a sample level. Specifically, for each sample, cells were excluded if they had fewer than 3,000 (or minus three MAD for internal samples) or more than the median plus three MAD of fragments overlapping peaks. Similar criteria were applied to the other QC metrics, with cells excluded for having FRiP scores below the median minus three MAD, blacklist fractions or nucleosome signals above the median plus three MAD, and TSS enrichment scores below the median minus three MAD.

While annotating broad and granular cell types and states on the integrated dataset, we further defined a set of cells to be excluded in downstream analysis. We excluded cells that appeared to be contaminants, have ambiguous cell identities, or clustered solely by a QC metric. For example, a contaminant cluster in the T cell compartment would be a cluster of cells with unexpectedly high accessibility of myeloid markers, indicating high ambient DNA or other artifacts. After noticing FRiP driving some clustering results, we set an additional sample-level cutoff on FRiP to remove cells below the median minus 1.25 MAD from downstream analyses.

### snATAC-seq data integration

To combine all QC-passing cells, we quantified all cells on a shared peak set. We used a peak set created from broad cell-type-level peak calling on the internal scATAC-seq only samples. We then constructed and combined these individual count matrices. Next, we performed normalization and dimension reduction using Signac RunTFIDF and RunSVD commands to perform latent semantic indexing (LSI). Then, we batch corrected the LSI components using Harmony with sample ID as the batch covariate ^21^. Using these batch-corrected embeddings, we constructed a nearest neighbor graph and performed clustering using the smart local moving (SLM) algorithm with the Seurat FindClusters function. Using gene activity scores and CopyscAT copy number estimates, we defined broad cell types and re-called peaks splitting by broad cell type ^48^. For all downstream analyses, we used this final broad cell type-specific peak set, and again constructed and combined count matrices, performed normalization, dimension reduction, and batch-corrected the LSI embeddings with Harmony.

### Clustering and cell type annotation

With the integrated dataset, we used the 2nd through 30th Harmony-adjusted LSI components to perform clustering with Seurat FindClusters using the SLM algorithm with a 0.1 resolution. We annotated these clusters with cell lineages (e.g., lymphoid, myeloid, stromal, tumor) using differential gene activity score analysis, inspection of insertion events over known marker genes, and copy number estimates. Per-cell copy number estimates were obtained using CopyscAT with default parameters ^48^. To annotate more granular cell types and states, we iteratively subsetted cells within each lineage and cell type, re-normalized with RunTFIDF, reduced dimensions with RunSVD, and clustered with the SLM algorithm using a range of resolutions and number of LSI components in Signac. We used a combination of differential gene activity score analysis, TF binding site motif enrichment, and copy number estimates to annotate groups of cells with distinct chromatin accessibility patterns as a particular cell type or state. Gene activity scores were computed using the GeneActivity function in Signac.

### Mutational data

For the internal cohort, where available, additional tissue was used for whole exome sequencing (WES) or panel sequencing via OncoPanel. For WES data, reads were aligned to hg19 reference using BWA v.0.5.9 ^49^. We utilized the Getz Lab CGA WES Characterization pipeline (https://portal.firecloud.org/#methods/getzlab/CGA_WES_Characterization_Pipeline_v0.1_Dec2018/) developed at the Broad Institute to call, filter and annotate somatic mutations and copy number variation. The pipeline employs the following tools: MuTect ^50^, ContEst ^51^, Strelka ^52^, Orientation Bias Filter ^53^, DeTiN ^54^, AllelicCapSeg ^55^, MAFPoNFilter^56^, RealignmentFilter, ABSOLUTE ^57^, GATK ^58^, PicardTools ^59,60^, Variant Effect Predictor ^61^, Oncotator ^62^.

Mutational data for the Wu et al cohort was obtained through the available Mutation Annotation Format (MAF) files stored in the Genomic Data Commons ^17^. We preferentially selected MAF files originating from the same sample ID as the snATAC-seq sample. If unavailable, we used mutations from the MAF file of another specimen from that patient. For Yu et al and Long et al cohorts, mutations were acquired from the studies’ respective supplemental tables ^19,20^.

Mutation data for the TCGA KIRC cohort was downloaded from cBioPortal (https://www.cbioportal.org/study/summary?id=kirc_tcga_pan_can_atlas_2018)^18,63–65^. WES data for the JAVELIN Renal 101 cohort were obtained from published results ^42,43^. Processing to retrieve mutation calls was performed the same as the internal cohort WES sequencing data as described above.

Putative loss-of-function (pLOF) mutations for all cohorts, including the internal cohort, Wu et al., Yu et al., Long et al., TCGA KIRC, and JAVELIN Renal 101, were defined as those with any of the following predicted consequences: frameshift variants (insertions or deletions), nonsense mutations (e.g., stop gained), nonstop mutations (loss of a stop codon), start lost, splice site mutations (e.g., splice donor or splice acceptor), or other truncating events.

### Clinical data

Clinical data for the Wu et al and Yu et al cohorts was obtained from the studies’ respective supplemental tables. Clinical data for the Long et al cohort was obtained from the studies’ supplemental table along with information from the study’s authors. All snATAC-seq cohorts’ mutational and clinical data relevant to the study is summarized in Table S1.

Clinical data for TCGA KIRC was downloaded from cBioPortal (https://www.cbioportal.org/study/summary?id=kirc_tcga_pan_can_atlas_2018)^18,63–65^.

Clinical data for JAVELIN Renal 101 was acquired from the published results ^42,43^.

### Differential gene accessibility

We identified genes with differential accessibility across non-malignant broad cell types using the logistic regression (LR) framework in Seurat’s FindMarkers function. This analysis included genes with accessibility in at least 10% of cells within each cell type and was adjusted for technical variability in sequencing depth. Additionally, we restricted analysis to a maximum of 1,000 cells per cell type and focused on genes with a minimum log-fold change of 0.1.

Similarly, we identified differentially accessible genes across RCC malignant cell states using the LR method, with accessibility in at least 20% of cells per state and a minimum log-fold change of 0.1, adjusting for sequencing depth across cells.

### Differential peak accessibility and peak sets

We identified differentially accessible peaks across ccRCC tumor states using the LR framework in Seurat’s FindMarkers function. This analysis focused on peaks with accessibility in at least 20% of cells per tumor state, adjusting for sequencing depth, and was limited to a maximum of 1,000 cells per state. Peaks were considered state-specific if they showed significantly higher accessibility in a given tumor state compared to all other states, with significance defined by an adjusted p-value below 0.05 (Bonferroni correction).

Similarly, we identified differentially accessible peaks between tumor cells from pLOF *BAP1*-mutated lesions and *BAP1* wild type lesions using the LR framework in Seurat’s FindMarkers function. We restricted the analysis to cells from ccRCC tumor states (C0,C1,C2,C3) from tumors with a pLOF mutation in *BAP1* or no non-silent mutations in *BAP1* (e.g., not including cells from samples with *BAP1* missense mutations). Peaks were considered enriched in *BAP1* pLOF or *BAP1* wild-type lesions if they were accessible in at least 10% of cells within a group, with an adjusted p-value below 0.05 (Bonferroni correction) and an absolute log2 fold-change greater than one.

### Peak set scoring

We scored cells for a given significant peak set by first computing the average normalized counts for the peak set. To do this, we summed the normalized counts over the peaks for each cell from the data slot. Then, we divided this value per cell by the number of peaks in the peak set.

Next, we constructed a reference peak set by finding peaks open in at least 10% of cells using the Signac AccessiblePeaks function. Then, we subsetted those peaks to ones that matched the significant peak set GC percent distribution using the Signac MatchRegionStats function. With this reference peak set, we computed the average normalized peak counts per cell in the same way as the significant peak set. Then, to get the final score per cell for a significant peak set we subtracted the reference peak set score from the significant peak set score.

### Transcription factor analysis

We identified TFs whose binding site motif was enriched in peak sets using the Signac FindMotifs function with default parameters which employs a hypergeometric test with FDR correction. As background peaks to provide to the function, we selected peaks that were accessible in at least 10% of all cells, where all cells are the total number of cells in all comparison groups for the given analysis. These peaks were further limited to those that match the distribution of GC content using the Signac MatchRegionStats function with default parameters.

### GREAT pathway analysis

We used the previously defined peak sets and converted them to hg19 coordinates using the package rtracklayer’s liftover function. Then, we retrieved enriched pathways per peak set using GREAT version 3.0.0 with default parameters ^22^.

### Data analysis and visualization

Most data analysis and visualization were performed with Signac version 1.10.0 unless otherwise specified ^66^.

### Derivation of epigenetic measure of interferon signaling

To derive an epigenetic measure of IFN signaling, we leveraged four tumor samples with both RNA and ATAC profiled from the same cells. First, we limited this dataset to putative tumor cells, as determined via the ATAC data, whose RNA and ATAC assays both passed QC. The ATAC-based QC was performed as described above. For RNA samples, we first performed ambient RNA decontamination on each sample using the remove-background module of CellBender (v0.2.0) with 150 training epochs, 25,000 total droplets included, and the “full” model ^67^. For each sample, the expected cells parameter was taken from the estimated number of cells from Cell Ranger. All other parameters were kept as default. Next, we performed multiplet detection and filtering on CellBender-cleaned matrices using the Scrublet package (Python, v0.2.1) with the expected doublet rate parameter set to 0.1 ^68^. We manually selected doublet score thresholds for each sample and removed predicted doublets, yielding singlet-only, ambient RNA-decontaminated counts matrices. Lastly, we removed cells with 200 or fewer genes detected or 5% or more of counts from mitochondrially encoded transcripts. We then normalized the raw counts using the default Seurat log-normalization method.

We used CytoSig to measure IFN1 and IFNG cytokine signaling from the transcriptomic data of these tumor cells ^69^. We supplied CytoSig with the raw RNA counts matrix generated using the DropletUtils v1.23.0 function write10xCounts on the Seurat object.

To determine which genomic regions’ accessibility were associated with the transcriptomic measures of IFN1 and IFNG signaling, we used a mixed effects modeling approach with the lmer function from the lme4 v1.1-32 R package. We only evaluated peaks that were called in tumor cells and were accessible in greater than 5% of the multiome tumor cells (n=80,768 peaks). For each of these genomic regions, we fit a mixed effects model for predicting accessibility using IFN1 or IFNG CytoSig score (from the transcriptomic data), sequencing depth, and the sample ID as predictor variables. In these models, the sample ID was considered a random effect, and all other variables were considered fixed effects. Peaks were considered associated with IFN1 or IFNG signaling if the IFN1 or IFNG β coefficient was greater than 0 and had a nominal p-value less than 0.05.

We used these peak sets to measure IFN signaling by following the peak set scoring methods described above, barring a requirement that the reference peaks be accessible in at least 5% of cells.

### Quantification of transposable element accessibility

With the position-sorted BAM output of the CellRanger ATAC pipeline, we first sorted this BAM by read name using Picard’s SortSam tool. We then utilized an adapted scTE method (available at https://github.com/sabrinacamp2/scTE) to assess TE accessibility in ccRCC tumor biopsies. This adaptation involved defining a maximum fragment length for paired-end alignments used in the TE quantification. With this adapted scTE method, we used the following command to produce a TE counts matrix: ‘scTEATAC -CB CB - UMI False -p 6 -i ${bam} --expect-cells 20000 -x ../hg38.te.atac.idx -o ${sample_id}’.

To generate a UMAP based on TE counts in Seurat, we first filtered to cells that passed our ATAC QC as described above. Then, we log normalized the TE counts with the scale factor being the median of the number of TEs detected per cell. We scaled the data and ran PCA. Using the top 15 PCs, we computed a nearest neighbors graph and computed a UMAP. Then, we colored each cell by its broad cell type annotation as determined from the peak (ATAC)-based analysis.

### Bulk RNA-seq data acquisition and alignment

Hg-38 aligned RNA-seq BAMs were downloaded from GDC for the TCGA KIRC cohort.

JAVELIN Renal 101 RNA-seq data were obtained from published results ^42,43^. These paired-end bulk RNA-seq samples were realigned to the hg19 genome reference using STAR v2.7.068, with alignment parameters --outFilterMultimapNmax 20 -- outFilterMismatchNmax 999 --outFilterMismatchNoverReadLmax 0.04 –alignIntronMin 20 --alignIntronMax 1250000 --alignMatesGapMax 1250000.

### Quantification and normalization of transposable element expression

We utilized the adapted scTE method, as described above, to quantify TE expression in these samples. We used the following command to produce a TE counts matrix: scTE -I ${bam} -o ${sample_id} -x /scTE/${genome}.exclusive.idx -CB False -UMI False --hdf5 False. The genome index was selected to match the reference genome that the RNA-seq data was aligned to. For the TCGA KIRC dataset, we used the hg38 genome index. For the JAVELIN Renal 101 dataset, we used the hg19 genome index. We normalized the expression data using DESeq v1.38.3 size factor normalization and variance stabilizing transformation.

### Linear regression model for BAP1 mutation and ERV3-16A3_LTR expression TCGA KIRC

To assess the relationship between ERV3-16A3_LTR expression and *BAP1* mutation status, we applied a linear regression model using the lm function from the base R stats package. Our analysis included only samples with a *BAP1* pLOF mutation or without any *BAP1* mutation (i.e. samples with non-truncating *BAP1* mutations were excluded). We further restricted to one sample per patient. In the model, we used normalized ERV3-16A3_LTR expression as the outcome variable and included predictor variables *BAP1* mutation status, tumor purity, disease stage, patient age at diagnosis, sex, and history of neoadjuvant treatment. Tumor purity and patient age were scaled to account for their continuous nature. Tumor purity is a scaled consensus tumor purity estimate from Aran et al ^70^. Additionally, we conducted stage-stratified analyses by separating the dataset into early-stage (stage I and II) and late-stage (stage III and IV) tumors.

### JAVELIN Renal 101

To evaluate the association between ERV3-16A3_LTR expression and *BAP1* mutation status, we fit a linear regression model with normalized ERV3-16A3_LTR expression as the outcome variable using the lm function from the base R stats package. For this analysis, we included samples that had complete RNA expression data, mutation information, FACETS-derived tumor purity estimates ^71^, and non-null values across key clinical variables (e.g., treatment arm, progression-free survival, overall survival, MSKCC risk score, sex, age, PD-L1 status). Only tumors with either a pLOF *BAP1* mutation or no *BAP1* mutation were considered. The model included *BAP1* mutation status, tumor purity, patient age at diagnosis, sex, and tumor location as predictors.

Tumor purity and patient age were scaled to adjust for their continuous nature. Tumor location specified whether the sample was from a primary or metastatic site.

### Survival models for ERV3-16A3_LTR expression TCGA KIRC

Sample inclusion criteria was the same as described above for TCGA KIRC linear regression model.

To assess the relationship between ERV3-16A3_LTR expression and overall survival, we employed a Cox proportional hazards model with overall survival time as the outcome, using the survival v3.5-5 package in R. OS served as the outcome variable. Predictor variables included ERV3-16A3_LTR expression, *BAP1* mutation status, tumor purity, disease stage, patient age at diagnosis, sex, and history of neoadjuvant treatment. Tumor purity and age were scaled to normalize continuous variables.

Additionally, we conducted stage-stratified analyses by separating the dataset into early-stage (stage I and II) and late-stage (stage III and IV) tumors.

### JAVELIN Renal 101

Sample inclusion criteria was the same as described above for JAVELIN Renal 101 linear regression model. To assess the relationship between ERV3-16A3_LTR expression and progression-free survival (PFS) in both the avelumab-plus-axitinib and sunitinib treatment arms, we used a Cox proportional hazards model with the survival v3.5-5 package in R. In this model, PFS served as the outcome variable. Predictor variables included ERV3-16A3_LTR expression, *BAP1* mutation status, tumor purity, patient age at diagnosis, sex, tumor location, MSKCC risk group, and PD-L1 positivity status as defined in the original publication. Tumor purity and age were scaled to normalize continuous data.

### Immunogenic ERVs

We limited the analysis to only ccRCC tumor cells that were assigned to one of the shared epigenetic states (C0,C1,C2,C3) and had quantified TE accessibility. These cells were scored for IFN1 and IFNG signaling using our IFN1 and IFNG peak sets and described peak set scoring method.

We restricted the TE features to ERVs by filtering for TE features containing ‘HERV’ or ‘ERV’ and including only those with non-zero accessibility in more than 10 cells, yielding a set of 58 ERVs.

For each ERV and each IFN epigenetic measure (IFN1 and IFNG), we applied a mixed effects model to predict IFN signaling based on ERV accessibility, sequencing depth, and sample ID using the lmer function from the lme4 v1.1-32 R package. In these models, sample ID was considered a random effect and all other variables fixed effects. ERVs were considered IFN-associated if they met both criteria: a q-value below 0.05 and a positive coefficient for both IFN1 and IFNG signaling measures.

### Code availability

Code for re-generating all major figures can be found at https://github.com/vanallenlab/snatac-rcc-manuscript.

---

**Figure S1.**
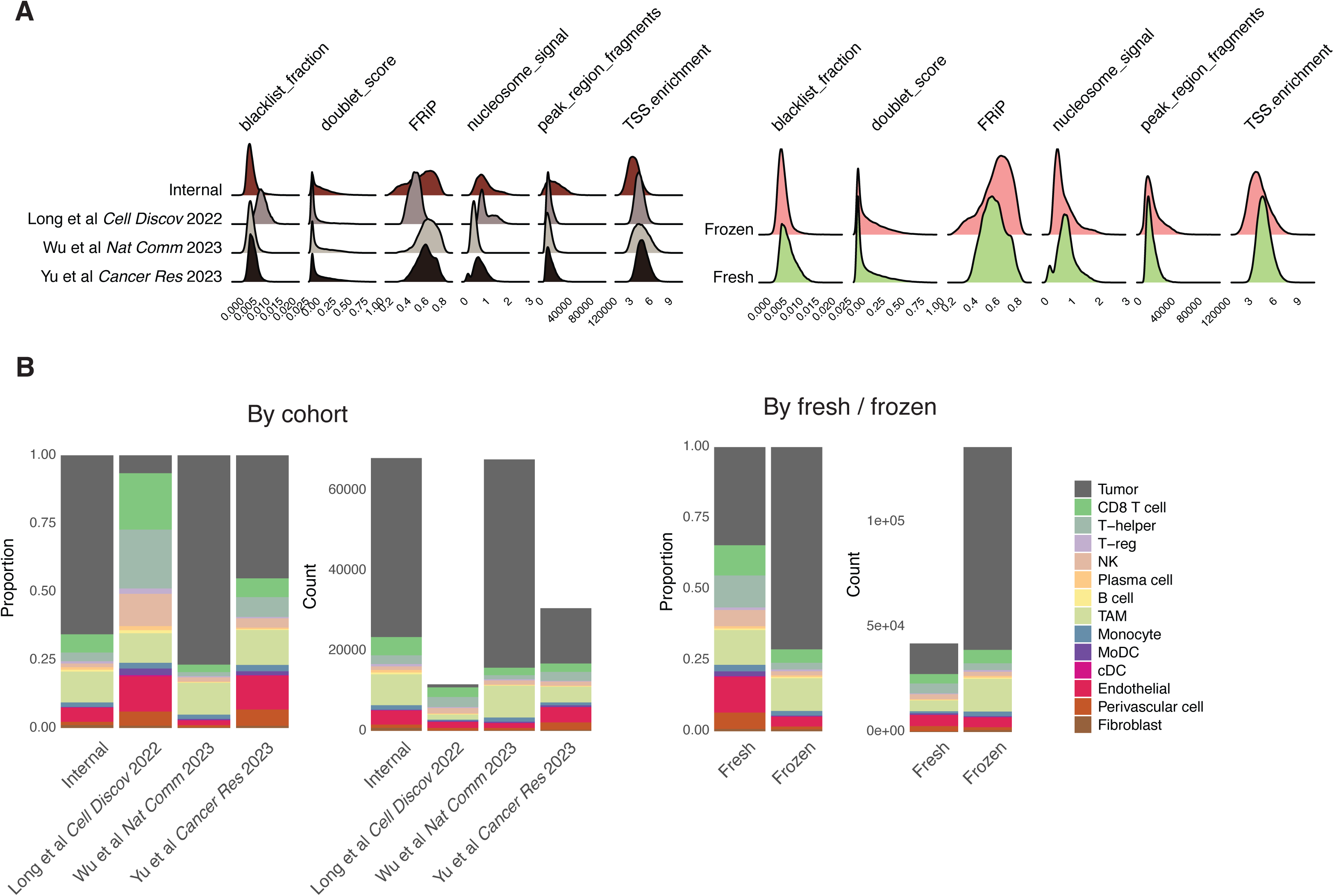
Quality control metric distributions and cell type capture. D) Density plots of quality control metrics split by cohort (left) and whether the sample was fresh or frozen (right) E) Proportional and count-based stacked bar plots detailing cell type capture per cohort (left) and whether the sample was fresh or frozen (right)

**Figure S2.**
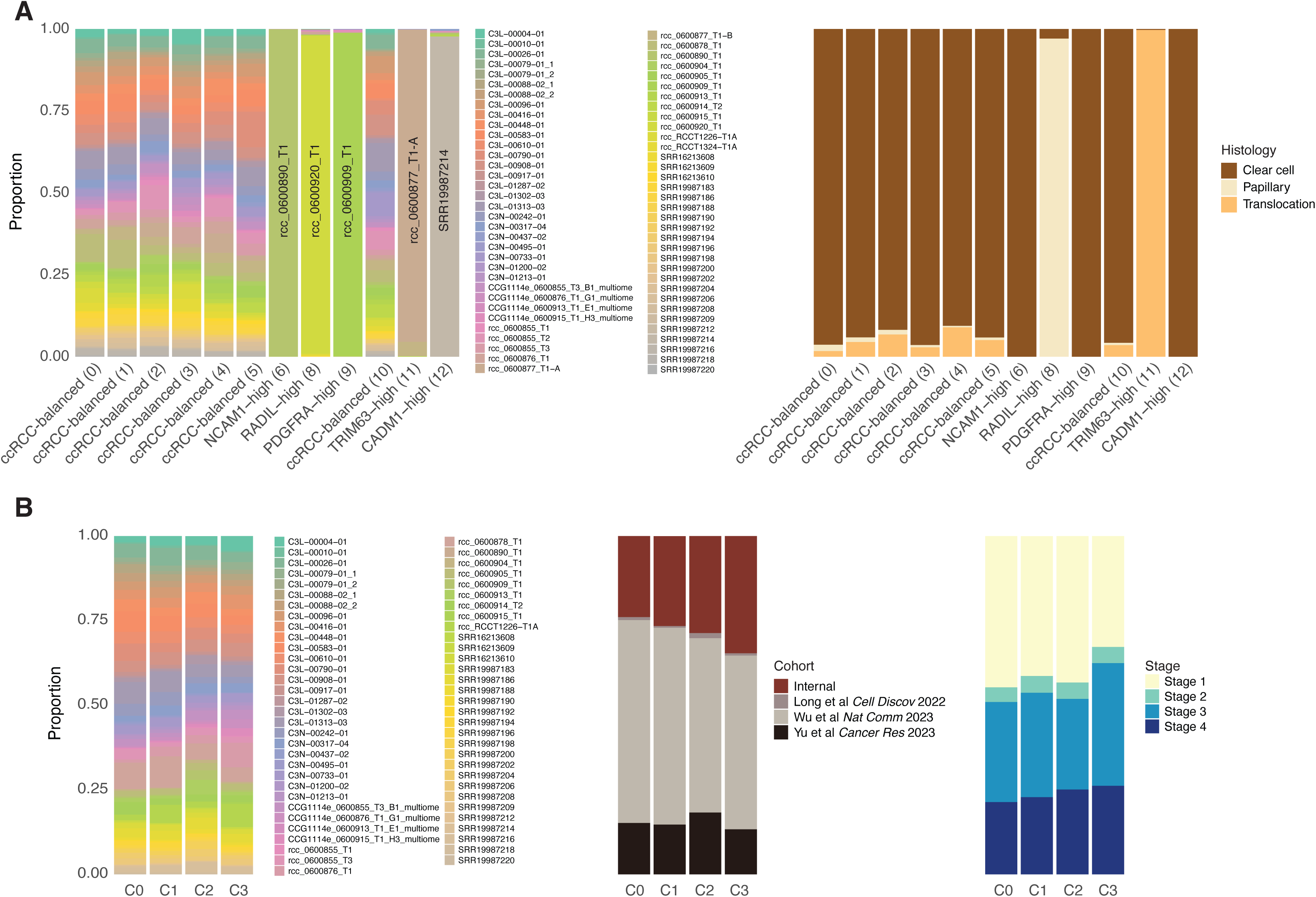
Feature proportions across RCC and ccRCC epigenetic states. A) Proportion of cells from each sample (left) and each histology (right) for each cluster identified in RCC tumor cells B) Proportion of cells from each sample (left), cohort (middle) and disease stage (right) for each cluster identified in ccRCC tumor cells

**Figure S3.**
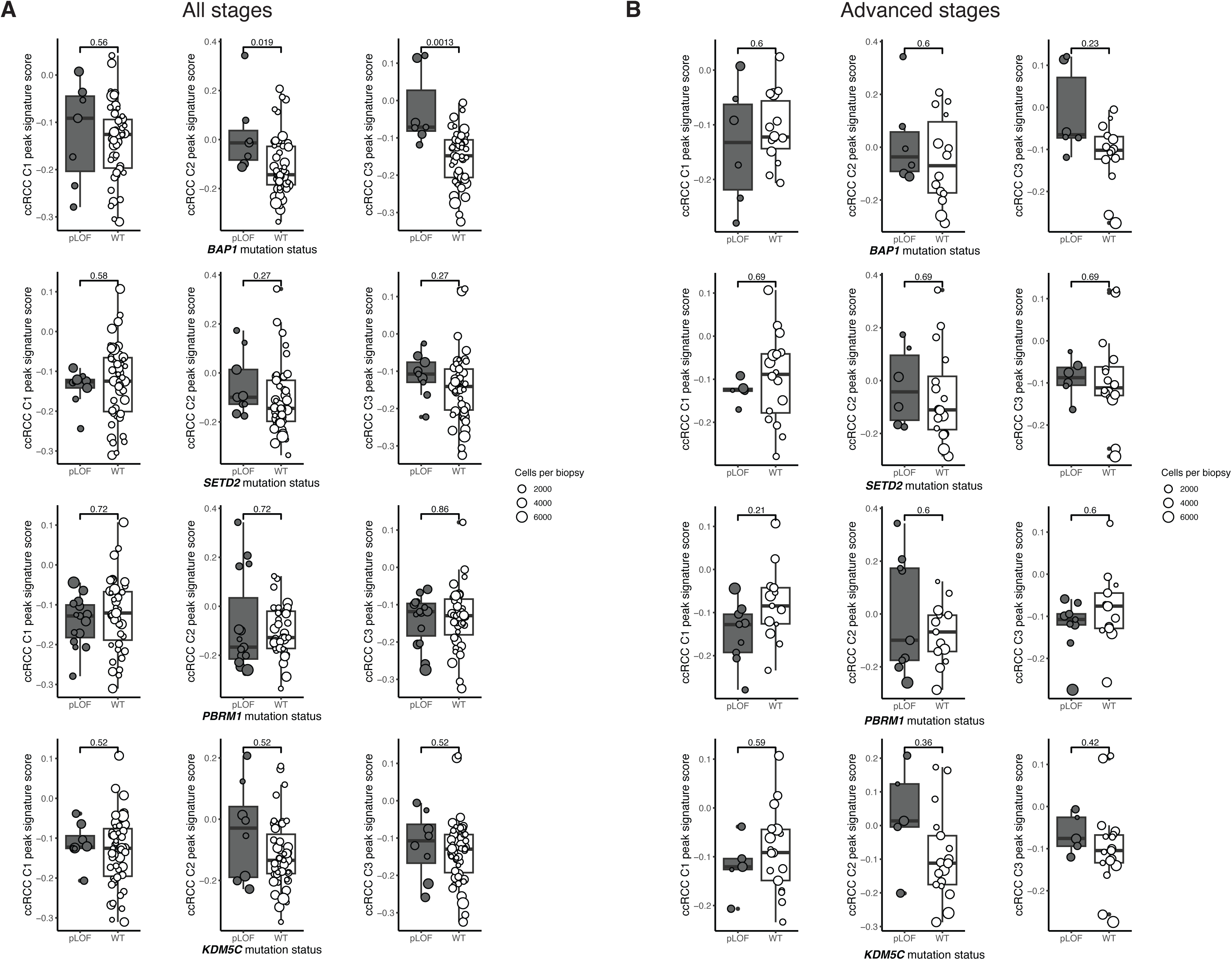
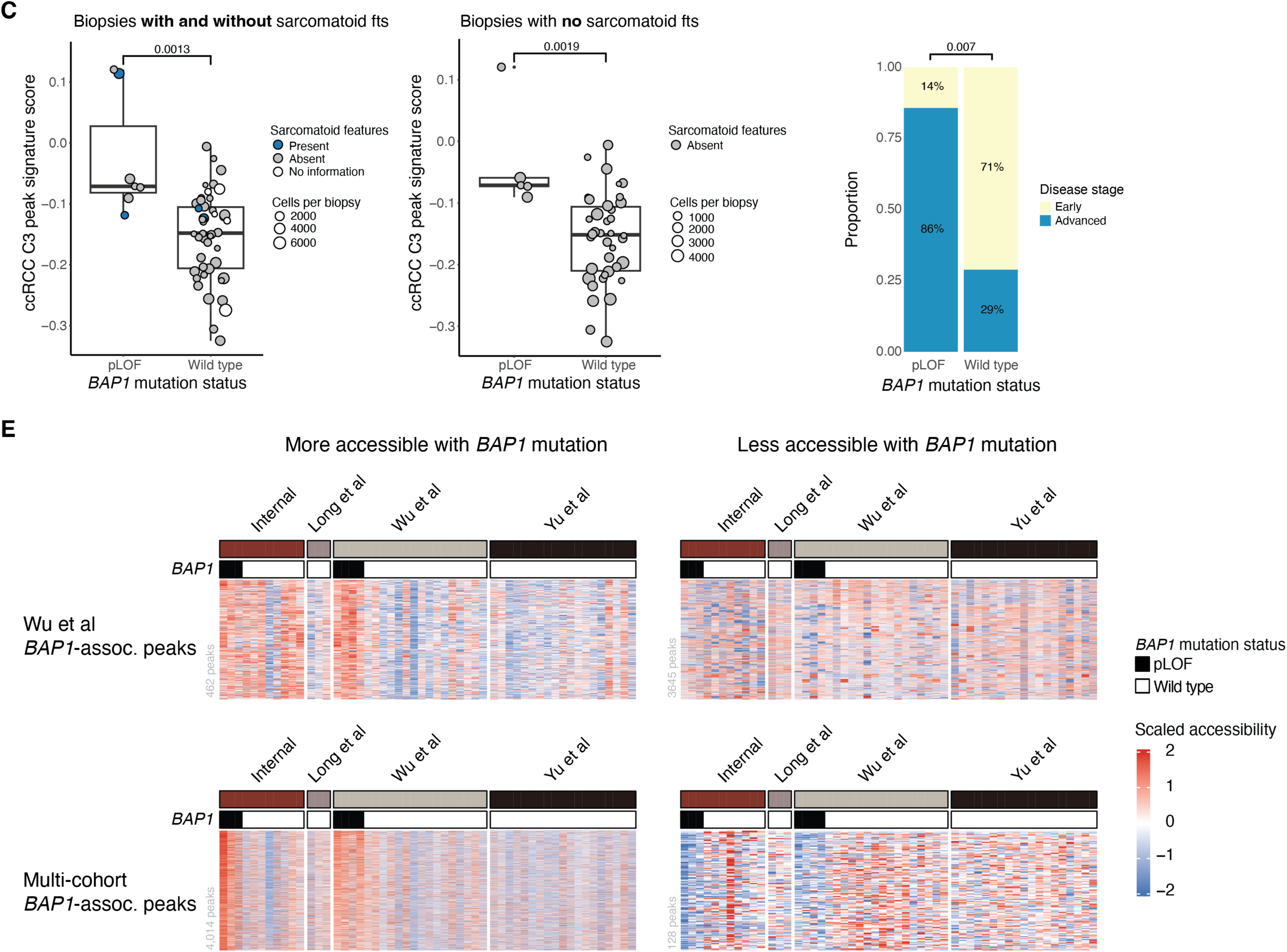
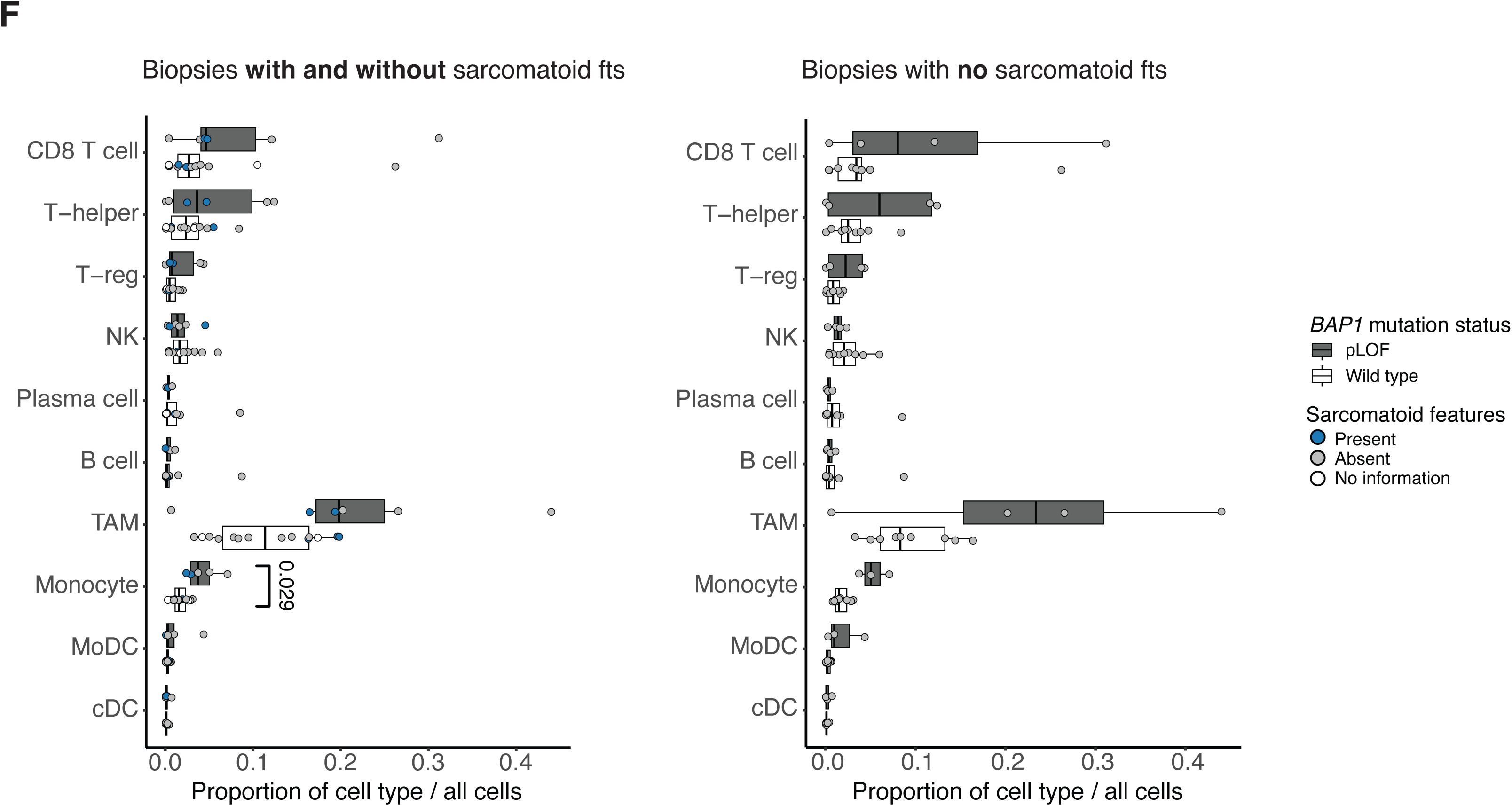
Epigenetic modifier mutations and ccRCC states, relationship between clinical variables and *BAP1* mutation and associated results, and evaluation of internal and external *BAP1* mutation associated peak sets. A) Boxplot of median cluster peak signature scores (C1, C2, C3 by column) per unique biopsy, split by epigenetic modifier mutation status (by rows). Wild type groups do not contain pLOF or non-silent mutations in the respective gene. Analysis limited to cells from ccRCC tumor states (C0,C1,C2,C3), Q-value shown for each comparison (two-sided Wilcoxen, FDR correction). Including biopsies from patients with all disease stages. B) Boxplot of median cluster peak signature scores (C1, C2, C3 by column) per unique biopsy, split by epigenetic modifier mutation status (by rows). Wild type groups do not contain pLOF or non-silent mutations in the respective gene. Analysis limited to cells from ccRCC tumor states (C0,C1,C2,C3). Q-value shown for each comparison (two-sided Wilcoxen, FDR correction). Including biopsies from patients with advanced disease stages. C) Boxplots of median ccRCC C3 peak signature score per unique biopsy, split by *BAP1* mutation status. *BAP1* wild type excludes pLOF and other non-silent mutations. Analysis limited to cells from ccRCC tumor states (C0,C1,C2,C3). Points colored by whether sarcomatoid features were annotated for the given biopsy. Left, including biopsies with or without sarcomatoid features annotated. Q-value displayed from original Figure 3A analysis (two-sided Wilcoxen, FDR correction). Right, only including biopsies that did not have sarcomatoid features. P-value displayed (two-sided Wilcoxen). D) Proportion of patients with early or advanced disease by *BAP1* mutation status. Two-sided Fisher’s exact p-value. E) Heatmaps of scaled peak accessibility scores for regions found to be associated with *BAP1* mutation in Wu et al (first row heatmaps) and in our multicohort late stage only analysis (second row heatmaps). The first column of heatmaps are peaks found to be significantly more accessible with a *BAP1* mutation in each analysis. The second column of heatmaps are peaks found to be significantly less accessible with a *BAP1* mutation in each analysis. Each column within one heatmap is a scaled average peak accessibility for a given biopsy. Columns are ordered by cohort and then *BAP1* mutation status. Analysis limited to cells from ccRCC tumor states (C0,C1,C2,C3). F) Boxplots comparing broad immune cell type proportions between biopsies with pLOF *BAP1* mutation and *BAP1* wild type, restricted to advanced stage disease patients. Points overlaying boxplots show per biopsy cell type proportions and are colored by whether sarcomatoid features were annotated for the given biopsy. Left, including biopsies with or without sarcomatoid features annotated. Q-value for monocyte proportion comparison shown (two-sided Wilcoxen, FDR correction); other cell type comparisons not significant. Right, only including biopsies that did not have sarcomatoid features.

**Figure S4.**
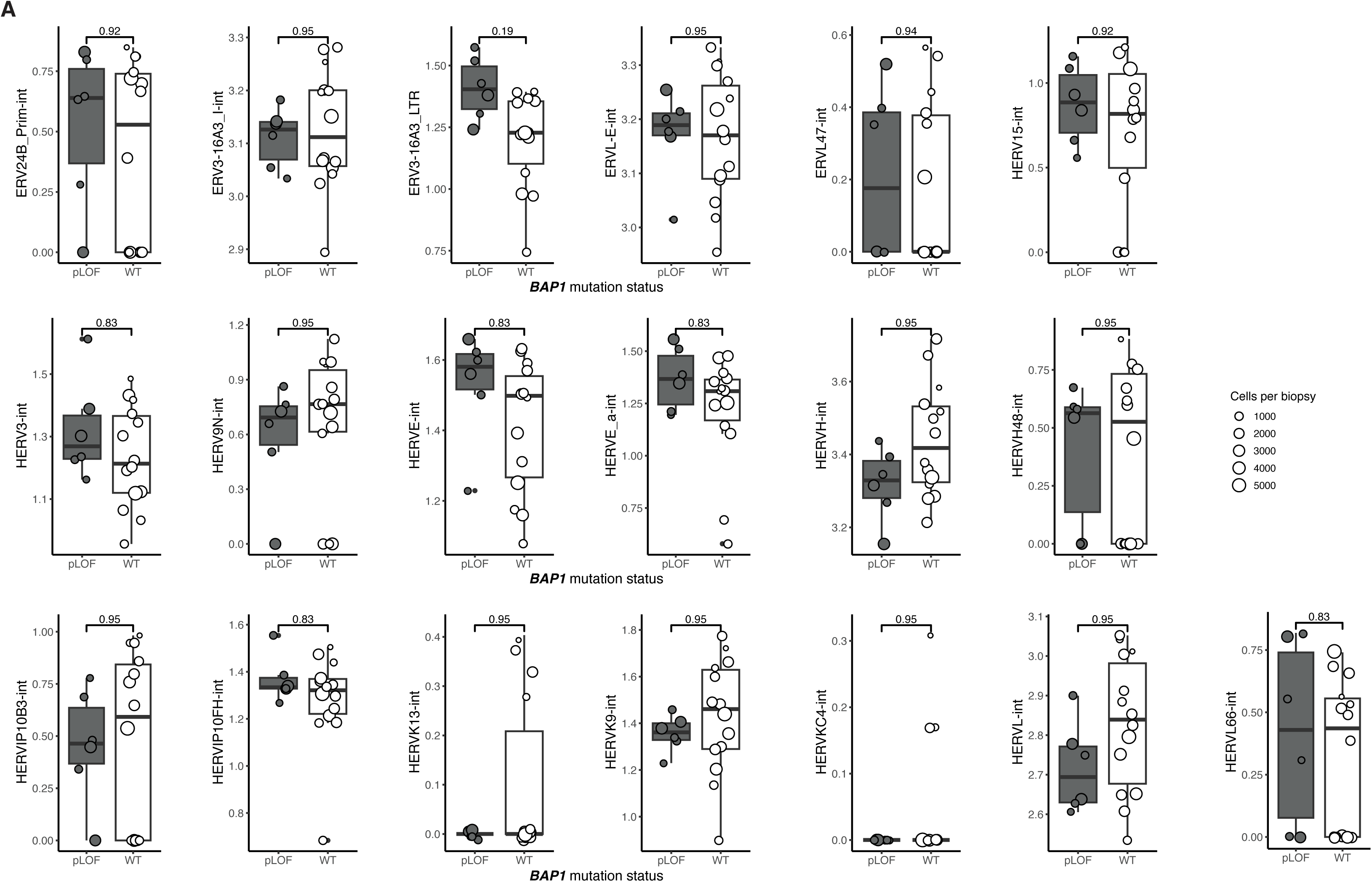
IFN-associated ERV accessibility and *BAP1* mutation status. A) Boxplot of median IFN-associated ERV accessibility per unique biopsy, split by *BAP1* mutation status. Wild type groups do not contain pLOF or non-silent mutations in *BAP1*. Analysis limited to cells from ccRCC tumor states (C0,C1,C2,C3). Q-values shown for each comparison (one-sided Wilcoxen, FDR correction). Including biopsies from patients with advanced disease stage

**Figure S5.**
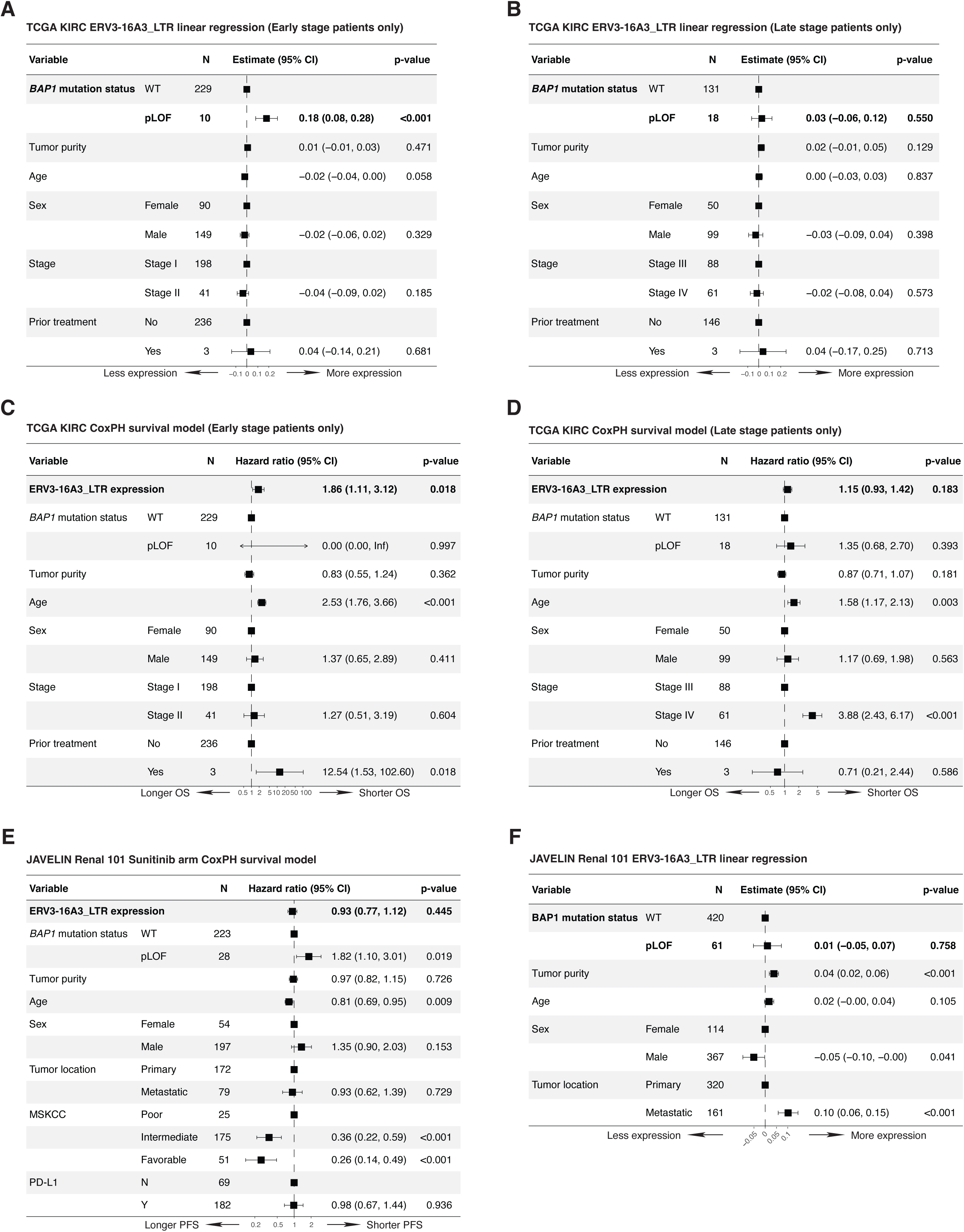
Disease stage-stratified TCGA KIRC analyses, Sunitinib-only arm survival model results in Javelin Renal 101, and relationship between ERV3-16A3_LTR expression and *BAP1* mutation status in Javelin Renal 101. A) Forest plot showing beta coefficients from a linear regression model for expression of ERV3-16A3_LTR in TCGA KIRC cohort. Restricted to patients with early stage disease. B) Forest plot showing beta coefficients from a linear regression model for expression of ERV3-16A3_LTR in TCGA KIRC cohort. Restricted to patients with late stage disease. C) Forest plot showing hazard ratios from a Cox proportional hazards model for overall survival in TCGA KIRC cohort. Restricted to patients with early stage disease. D) Forest plot showing hazard ratios from a Cox proportional hazards model for overall survival in TCGA KIRC cohort. Restricted to patients with late stage disease. E) Forest plot showing hazard ratios from a Cox proportional hazards model for progression-free survival in Sunitinib-only treatment arm of JAVELIN Renal 101 cohort. F) Forest plot showing beta coefficients from a linear regression model for expression of ERV3-16A3_LTR in both arms of JAVELIN Renal 101 cohort

